# Phylogeographic parallelism: concordance of patterns in closely related species illuminates underlying mechanisms in the historically glaciated Tasmanian landscape

**DOI:** 10.1101/548446

**Authors:** K. Kreger, B. Shaban, E. Wapstra, C.P. Burridge

**Affiliations:** Discipline of Biological Sciences, School of Natural Sciences, University of Tasmania, Private Bag 55, Hobart, Tasmania 7001, Australia; Australian Genomics Research Facility, Parkville, Victoria, Australia; Centre for System Genomics, University of Melbourne, Parkville, Victoria, Australia

**Author notes:** Ph, e-mail (CPB).

## Abstract

Phylogeography provides a means to understand mechanisms that shaped the distribution and abundance of species, including the role of past climate change. While concordant phylogeographic relationships across diverse taxa suggest shared underlying mechanisms (“phylogeographic parallelism”), it is also possible that similar patterns are the product of different mechanisms (“phylogeographic convergence”), reflecting variation among taxa in factors such as environmental tolerances, life histories, and vagility. Hence, phylogeographic concordance among closely related and ecologically similar species can yield a more confident understanding of the past mechanisms which shaped their distribution and abundance. This study documented mitochondrial and nuclear phylogeographic patterns in the ectotherm skink, *Niveoscincus metallicus*, which occupies historically glaciated regions of Tasmania, and contrasted these with the closely related and broadly sympatric *N. ocellatus*. Major phylogeographic breaks were similar in location between the two species, and indicative of isolation caused by retreat from high elevation areas during glaciations, but with long-term persistence at multiple low elevation sites. Hence, Pleistocene glacial refugia were altitudinal rather than latitudinal, a pattern mirrored in other temperate Southern Hemisphere taxa. This study also examined phylogeographic patterns across the intermittently inundated Bassian Isthmus between mainland Australia and the island of Tasmania, and revealed that structuring is similarly maintained when populations were physically isolated during interglacial rather than glacial stages.

## Introduction

Phylogeography provides a means to understand factors that shaped the distribution and abundance of species, including their responses to past climates (Hewitt, 2004). Past population dynamics are recorded in the genes of species, and robust quantitative frameworks are developing for the incorporation of such information into projections of species range and abundance (Fordham *et al.*, 2014), which is relevant for biodiversity conservation (Blois *et al.*, 2010). However, the value of phylogeography depends on the extent to which its results can be predicted across other regions and taxa (e.g., Bermingham & Avise, 1986; Avise, 1992; Carstens *et al.*, 2005; DeChaine & Martin, 2006). In this context, patterns from mid- to high-latitude regions of Western Europe and North America became paradigmatic within phylogeography (Taberlet *et al.*, 1998; Hewitt, 2000, 2004); elevated and endemic genetic variation at low latitude regions and lower genetic variation and phylogeographic structure across higher latitudes was interpreted as signatures of recolonization of previously glaciated areas from low latitude refugia. However, investigations in less extensively glaciated or unglaciated regions of the world often failed to recover patterns consistent with widespread extirpation and subsequent postglacial recolonisation from sparse refugia (e.g., Lessa *et al.*, 2003; Byrne, 2008; Byrne *et al.*, 2008; Byrne *et al.*, 2011; Qiu *et al.*, 2011; Sersic *et al.*, 2011; Breitman *et al.*, 2012; Lorenzen *et al.*, 2012; Turchetto-Zolet *et al.*, 2013). Other studies have also indicated long-term persistence with relatively subtle range movements through time in areas heavily affected by Pleistocene glaciations (Rull, 2009; Tzedakis *et al.*, 2013; de Lafontaine *et al.*, 2014). Therefore, phylogeographic patterns, and the inferences drawn from them, could be influenced by idiosyncrasies related to region or species (but these are also of value; see Papadopoulou & Knowles, 2016).

While concordant phylogeographic patterns are considered indicative of shared underlying histories, diverse phylogeographic responses to the same environmental change are expected among species owing to variation in environmental tolerances, life histories, and vagility (Jackson & Overpeck, 2000; Soltis *et al.*, 2006; Stewart *et al.*, 2010; Papadopoulou & Knowles, 2016). Therefore, even when shared patterns are observed among species, these may represent “phylogeographic convergence”—reflecting coincidence of patterns from different mechanisms—rather than “phylogeographic parallelism”, or the operation of the same mechanism. It is now widely advocated that studies attempting to attribute differences in phylogeographic patterns to differences in species attributes benefit from comparisons of closely related species, because these will differ in fewer other potentially confounding variables (Williams, 1994; Turner & Trexler, 1998; Dawson *et al.*, 2002; Whiteley *et al.*, 2004; Hickerson & Cunningham, 2005; Burridge *et al.*, 2008). The corollary of this is that shared patterns among closely related taxa are less likely to reflect phylogeographic convergence, and more likely to represent robust replicates for the inference of underlying mechanisms (Dawson, 2012; Dawson *et al.*, 2014).

Reptiles have been highlighted as excellent model species in phylogeographic studies (Camargo *et al.*, 2010). As ectotherms, they depend on heat from the environment to sustain metabolism and reproduction, and are therefore sensitive to changes in climate (Sinervo *et al.*, 2010). Reptiles also tend to have limited dispersal abilities (Metts *et al.*, 2000), which means that the genetic signatures of historical distributional changes are likely to persist for longer than in more vagile species such as large mammals and birds (Araújo *et al.*, 2008). Therefore, we expect that comparative phylogeographic studies of closely related reptiles will be more likely to observe shared patterns reflecting the same underlying mechanism, or where patterns differ, they can be more confidently ascribed to differences among these species with respect to morphology, physiology, life history, or behaviour. While many phylogeographic studies have exploited reptiles to increase our understanding of faunal responses to environmental change (Dubey & Shine, 2010; Breitman *et al.*, 2012; Vera-Escalona *et al.*, 2012; Nelson-Tunley *et al.*, 2016), few have analysed closely related species (e.g., Hare *et al.*, 2008).

There have been substantial efforts to overcome the early dearth of phylogeographic studies on Southern Hemisphere taxa (Beheregaray, 2008), and some generalisations have been made regarding biotic responses to Pleistocene climate fluctuations in this region, and contrasting with patterns in the Northern Hemisphere (Byrne, 2008; Byrne *et al.*, 2008; Wallis & Trewick, 2009; Byrne *et al.*, 2011; Sérsic *et al.*, 2011; Wallis *et al.*, 2016). Tasmania—the only Australian location that was directly influenced by Pleistocene glaciations (Colhoun & Barrows, 2011)—experienced Last Glacial Maximum (LGM) temperature depressions up to 6.5°C (Colhoun, 1985; Fletcher & Thomas, 2010), with tree-lines close to the modern day sea level (Gibson *et al.*, 1987). While Tasmania has been the subject of several phylogeographic studies (Chapple *et al.*, 2005; Macqueen *et al.*, 2009; Worth *et al.*, 2009; Zhang *et al.*, 2014) (McKinnon *et al.*, 2004; Nevill *et al.*, 2010; Worth *et al.*, 2011), it remains unclear whether its fauna responded to Pleistocene glaciations by widespread contraction into geographically discrete refugia, or instead persisted in multiple regions. Cliff *et al.* (2015) have conducted the most comprehensive range-wide phylogeographic study of a Tasmanian vertebrate, the spotted snow skink *Niveoscincus ocellatus*, and revealed regional persistence but also contrasting patterns throughout the species’ range: there was evidence of population persistence and limited gene flow within the historically and presently milder northeast, while the southeast and glacially-effected central regions were characterised by patterns consistent with greater post-glacial gene flow and demographic expansion.

We conducted a phylogeographic study in the metallic snow skink *Niveoscincus metallicus*, to contrast against the previous results for the congeneric *N. ocellatus* (Cliff *et al.*, 2015). Both species are widespread across Tasmania and occupy a broad elevational range, from sea level to ∼1000-1200 m (Figure 1), roughly equivalent to the regional subalpine tree line (Melville & Swain, 1999), and therefore have similar thermal niches (Melville & Swain, 2003; Caldwell *et al.*, 2017). *Niveoscincus metallicus* is also distributed across the islands of Bass Strait and into southeast mainland Australia (Figure 1). While *N. metallicus* is typically found in close proximity to deep litter or other shelter, such as fallen logs or rocks, *N. ocellatus* is obligate saxicolous (Melville & Swain, 1997). The analysis of *N. metallicus* enables us to test phylogeographic convergence with *N. ocellatus*, and if present, provide stronger inference of the mechanism underlying their phylogeographic history. We also assess the phylogeographic relationships of Tasmanian, mainland Australian, and Bass Strait *N. metallicus* populations with respect to their persistence and isolation across the periodically emergent Bassian Isthmus, a question relating to Pleistocene sea level change (Lambeck & Chappell, 2001) that has been addressed for a range of taxa (Chapple *et al.*, 2005; Keogh *et al.*, 2005; Symula *et al.*, 2008; Dubey & Shine, 2010; Chapple *et al.*, 2011b; Haines *et al.*, 2014; Ng *et al.*, 2014).

**Figure 1.**
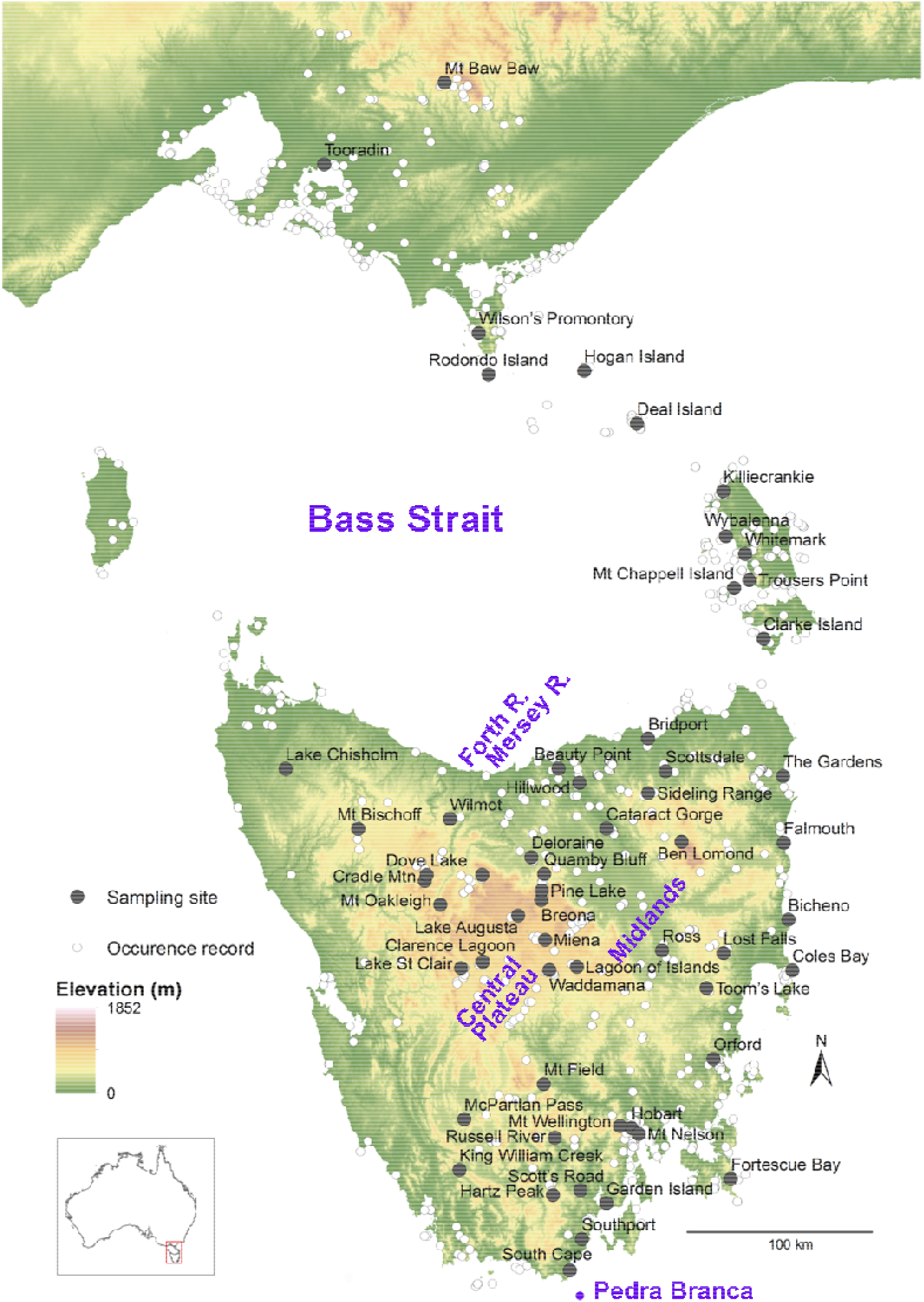
Recorded distribution of *Niveoscincus metallicus* (small hollow circles) and locations of sampling sites (dark grey circles). Base map: 250 m Digital Elevation Model (GeoScience Australia).

## Materials and Methods

Tissue was obtained from a total of 779 *N. metallicus* from 56 locations across the species’ geographic and elevational range (Figure 1, Supplementary Table 1), with the exception of King Island and southwest Tasmania. Sites were selected to ensure coverage of the same area investigated by Cliff *et al.* (2015) for *N. ocellatus*. Field collections were supplemented by institutional collections (Supplementary Table 2). Between 1–23 individuals were sampled per site (Supplementary Table 1). We employed *N. ocellatus* and *N. pretiosus* as outgroups. We sequenced several *N. pretiosus*, but also noted that other d assigned to this species (GenBank accession numbers HQ454789, DQ675234) fell deep within the *N. metallicus* tree, and we included these in our study as *N. metallicus*.

Genomic DNA extraction and PCR amplification of partial mitochondrial ND2 (*c.* 543 bp) and ND4 (*c.* 864 bp), and partial nuclear β-globin (intron 2, *c.* 782 bp) were conducted following Cliff *et al.* (2015). These regions exhibited useful levels of intraspecific genetic diversity in previous studies of squamates (Bell *et al.*, 2010; Chapple *et al.*, 2011b). Sequencing of both mtDNA strands was conducted in instances where sequence quality was low. All β-globin sequences were obtained from both strands. DNA sequences were edited and aligned in Geneious version 6.1.8 (Biomatters).

Ambiguous double peaks were observed in some ND2 sequence chromatograms, probably representing a nuclear pseudogene (Bensasson *et al.*, 2001). The protein coding regions of both mitochondrial genes were translated to ensure that sequences did not contain premature stop codons, that would indicate a divergent non-coding pseudogene. ND2 sequences from 300 individuals were obtained, and ND4 sequences were obtained from 340 individuals, representing 57 localities (Supplementary Table 1). Sequences from individuals where both ND2 and ND4 amplified successfully were concatenated, giving a total of 1406 bp for 284 individuals from 52 sites. A total of 72 individuals with sequence data for only one mitochondrial marker were omitted from the concatenated dataset, due to concerns about the effect of large amounts of missing data on phylogenetic inference (Lemmon *et al.*, 2009). Two sampling sites (Mount Bischoff and Cataract Gorge) were entirely omitted from the concatenated dataset because of substantial double peaks in ND2 sequences.

β-globin sequences were obtained from 84 individuals (168 alleles), representing 35 localities (Supplementary Table 1). Several individuals were excluded from this dataset based on heterozygous length mutations, resulting in long sections of superimposed chromatogram traces. PHASE 2.1.1 (Stephens *et al.*, 2001) was used to infer allelic states for individuals that were homozygous for sequence length but heterozygous at more than one nucleotide. PHASE input files were prepared and post-processed in SEQPHASE (Flot, 2010).

Bayesian phylogenetic reconstruction was conducted in MrBayes 3.2.2 (Ronquist *et al.*, 2012) on unique haplotypes identified from concatenated sequences. Individuals sharing the same sequence were assigned to haplotypes using TCSv1.21 (Clement *et al.*, 2000), ensuring that individuals with missing or ambiguous data at the same nucleotide position, but with otherwise identical sequences, were assigned to the same haplotype. mtDNA sequences were partitioned by gene region and analysed under substitution models suggested by jModelTest 2.1.7 (Guindon & Gascuel, 2003; Darriba *et al.*, 2012) using the Bayesian Information Criterion. When the most supported substitution model was not available in MrBayes, the next most complex model available was implemented. For ND4, TrN+I+Γ was selected by jModelTest and GTR+I+ Γ was implemented in MrBayes, and for ND2, TrN+Γ was selected and GTR+Γ was implemented. Two parallel runs were completed in MrBayes, each of 5.5 x 10^6^ generations, with four Markov Chain Monte Carlo (MCMC) chains with a heating parameter of 0.1 and sampling every 500 generations. The initial 25% of samples were discarded as burnin. Log files were checked using Tracer 1.6 (Rambaut & Drummond, 2013) to ensure that effective sample sizes were greater than 200 and that stationarity and convergence were achieved during model runs. The consensus tree was viewed and edited in FigTree v1.4.2 (Rambaut, 2014). Node support was assessed by posterior probabilities, with values greater than or equal to 0.95 considered to support the branching pattern.

Estimates of molecular diversity (haplotype diversity and nucleotide diversity) were calculated in Arlequin 3.5 (Excoffier & Lischer, 2010). The number of polymorphic and parsimony informative nucleotide sites and mean uncorrected genetic distances (p-distances) between sequences assigned to each mitochondrial clade were calculated in MEGA6.0 (Tamura *et al.*, 2013). Population structure was quantified and assessed for mitochondrial and nuclear data using two approaches: Analysis of Molecular Variance (AMOVA; Excoffier *et al.*, 1992) and Spatial Analysis of Molecular Variance (SAMOVA; Dupanloup *et al.*, 2002). AMOVA was performed in Arlequin. Sequence variation was hierarchically partitioned based on the membership of individuals to (i) major regional mitochondrial clades (Φ_CT_; clades defined from Bayesian phylogenetic analysis), (ii) geographic sites within clades (Φ_SC_), and (iii) geographic sites across the entire study range (Φ_ST_). The significance of estimated fixation indices was tested using 1000 permutations of the data. SAMOVA was performed using K = 2 to 7 groups in the SAMOVA package.

Divergence dating analyses were completed in BEAST 2.2.1 (Bouckaert *et al.*, 2014). Sequences were partitioned by gene region and substitution models suggested by jModelTest were implemented. To test for significant variation in mutation rates among different branches, a relaxed lognormal molecular clock model was implemented with a coalescent constant population size prior, run for 50 x 10^6^ generations and sampled every 5000 generations. Given that the rate variation parameter was close to zero, indicating that rates did not vary significantly between branches, a strict molecular clock was employed for estimating divergence times (Drummond & Suchard, 2010) with a normally distributed clock prior of mean 1.52% (standard deviation 0.5%) sequence divergence per million years (which equates to a lineage rate of 0.0076 substitutions per site per million years with a standard deviation of 0.0025). This rate was derived from several previous estimates of mitochondrial divergence in squamates based on fossil calibrations (Chapple *et al.*, 2011b). Results were visualised in Tracer to ensure that effective sample sizes were greater than 200 and that model runs had converged. The first 25% of samples were discarded as burnin and TreeAnnotator (Bouckaert *et al.*, 2014) was used to produce a maximum clade credibility tree with mean node heights. Uncertainty in node ages was assessed by examining the 95% high posterior density of values.

Due to shallow levels of divergence and the potential for reticulate evolution in nuclear sequences (Huson & Bryant, 2006), allele networks rather than phylogenetic trees were reconstructed for β-globin using the TCS statistical parsimony algorithm (Clement *et al.*, 2000), implemented in PopART (http://popart.otago.ac.nz/index.shtml; Leigh & Bryant, 2015). Congruence of relationships among alleles and each individual’s assignment to a mitochondrial clade were assessed visually. Alignments of phased sequences revealed several insertions or deletions (indels), which were present in a homozygous condition within a particular individual. Indels were coded as binary characters using SeqPhase (Muller, 2006). Network analyses with and without indel characters did not differ in topology and further analyses omitted indel information.

## Results

### Mitochondrial variation

Mitochondrial sequences were highly variable; from a total of 1406 base pairs, 439 sites were polymorphic and 304 sites were parsimony informative. A total of 215 unique haplotypes were identified from 284 individuals with sequence data for both ND2 and ND4. Overall haplotype diversity was high (H = 0.99, Table 1), with few haplotypes represented by more than one individual or more than one locality.

**Table 1.** Genetic diversity quantified by haplotype diversity (H) and nucleotide diversity (π) among groups and all samples based on 1406 base pairs of ND2 and ND4 mitochondrial sequence for *Niveoscincus metallicus*. Individuals were allocated to groups defined by regional mitochondrial clades. Sequences from admixed sites representing two mitochondrial clades were included in the analysis.

| Clade / subclade | H | $\pi$ |
| --- | --- | --- |
| SE | $0.9974 \pm 0.0016$ | $0.01608 \pm 0.00789$ |
| NE | $0.9938 \pm 0.0030$ | $0.02038 \pm 0.00997$ |
| - mainland Australia | $0.9692 \pm 0.0196$ | $0.00046 \pm 0.00024$ |
| - Bass Strait | $0.9715 \pm 0.0196$ | $0.02014 \pm 0.01013$ |
| - NE Tasmania | $0.9905 \pm 0.0114$ | $0.00903 \pm 0.00463$ |
| NW | $0.9942 \pm 0.0193$ | $0.01229 \pm 0.00645$ |
| S | $0.9841 \pm 0.0107$ | $0.01147 \pm 0.00582$ |
| Total | $0.9985 \pm 0.0005$ | $0.03985 \pm 0.01911$ |

Bayesian phylogenetic analysis of mitochondrial haplotypes recovered four major clades: a northwestern, a northeastern (including mainland Australia and the Bass Strait islands), a southeastern and a southern clade (Figure 2). These clades were geographically distinct, highly divergent and received strong posterior support (Figure 2). The level of phylogeographic structure varied within the major clades, with the northeast clade the most geographically structured (Figure 2). Northeast Tasmania was monophyletic, as were mainland Australia and each of the Bass Strait islands (with the exception of Flinders Island and the adjacent Mt Chappell Island), with strong posterior support (Figure 2). The order of divergence between these clades was well resolved with the exception of Clarke Island (Figure 2). The mainland Australian clade was nested among the Bass Strait island clades. There was relatively little phylogeographic structure among populations from the northeast of Tasmania. The southeast clade encompassed the largest land area, and several lineages were geographically clustered although not monophyletic (e.g. the northern Midlands, the east coast, Mt Wellington and surrounds, the eastern Central Plateau and the western Central Plateau; Figure 2). The northwest clade was the most divergent; however, this was the least internally structured geographically (Figure 2). The southern clade had limited evidence of spatial structuring (Figure 2). Geographic overlap between southeast and northwest clades occurred at Lake St Clair, while overlap between southeast and northeast clades occurred in northeast Tasmania at Hillwood and Ben Lomond, each representing high elevation sites (Figure 3). Uncorrected genetic distances between major clades were considerable, with approximately 7% sequence divergence separating the northwest clade, while the remaining clades were separated from one another by 4–5% sequence divergence.

**Figure 2.**
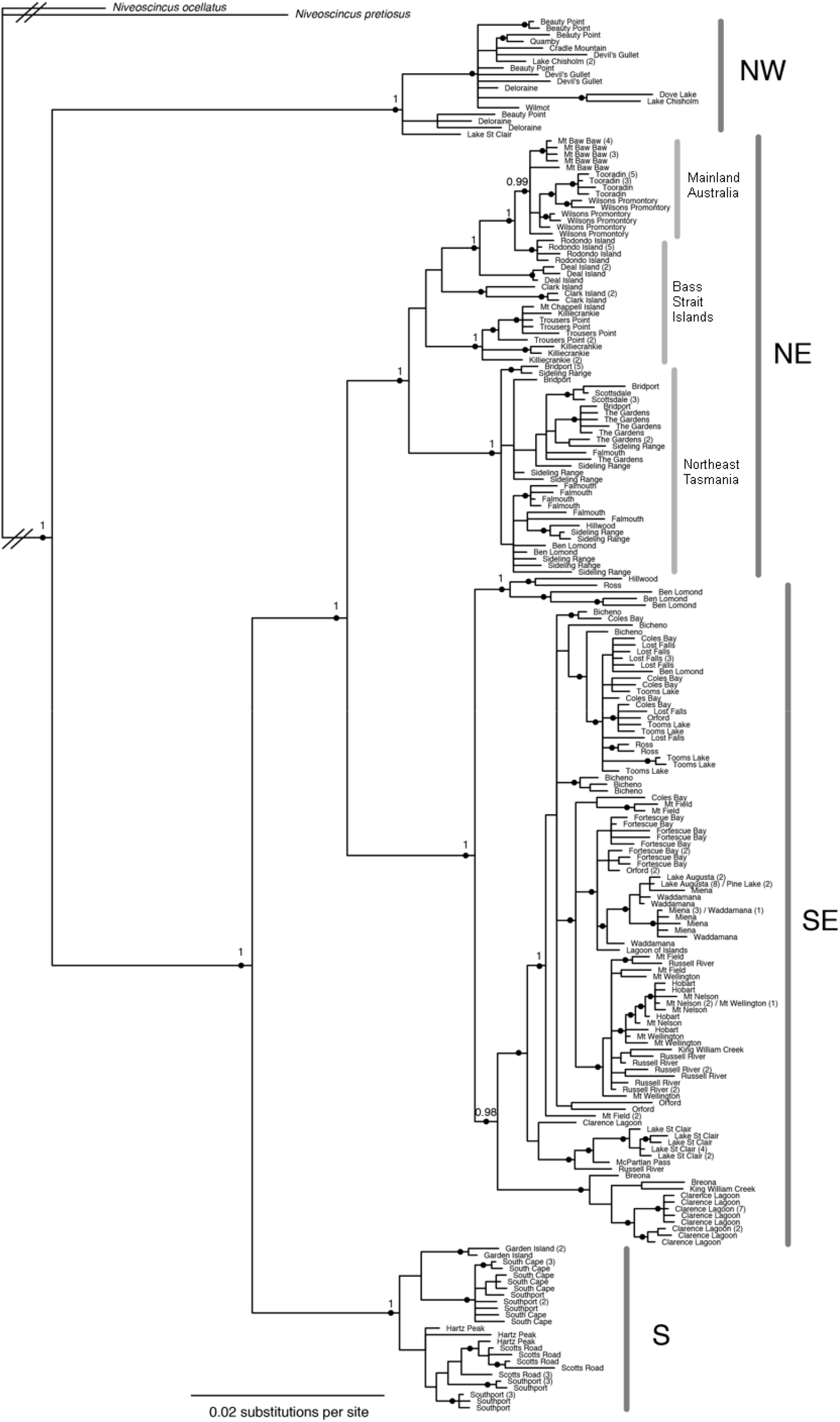
Bayesian inference tree of *Niveoscincus metallicus* reconstructed from 1406 bp of ND2 and ND4 mitochondrial DNA sequence. Major regional clades and major subdivisions within clades are indicated by vertical grey bars. Posterior probabilities >0.95 are indicated by black dots. Posterior probability values are included above key nodes. Branch lengths are scaled proportional to the scale bar. Branches to outgroups were truncated (indicated by dashed lines across branches). Analysis was conducted on unique haplotypes, and the numbers in parentheses after site names indicate the number of individuals belonging to that haplotype at that site. If only one individual per site was recorded, numbers in parentheses were omitted.

**Figure 3.**
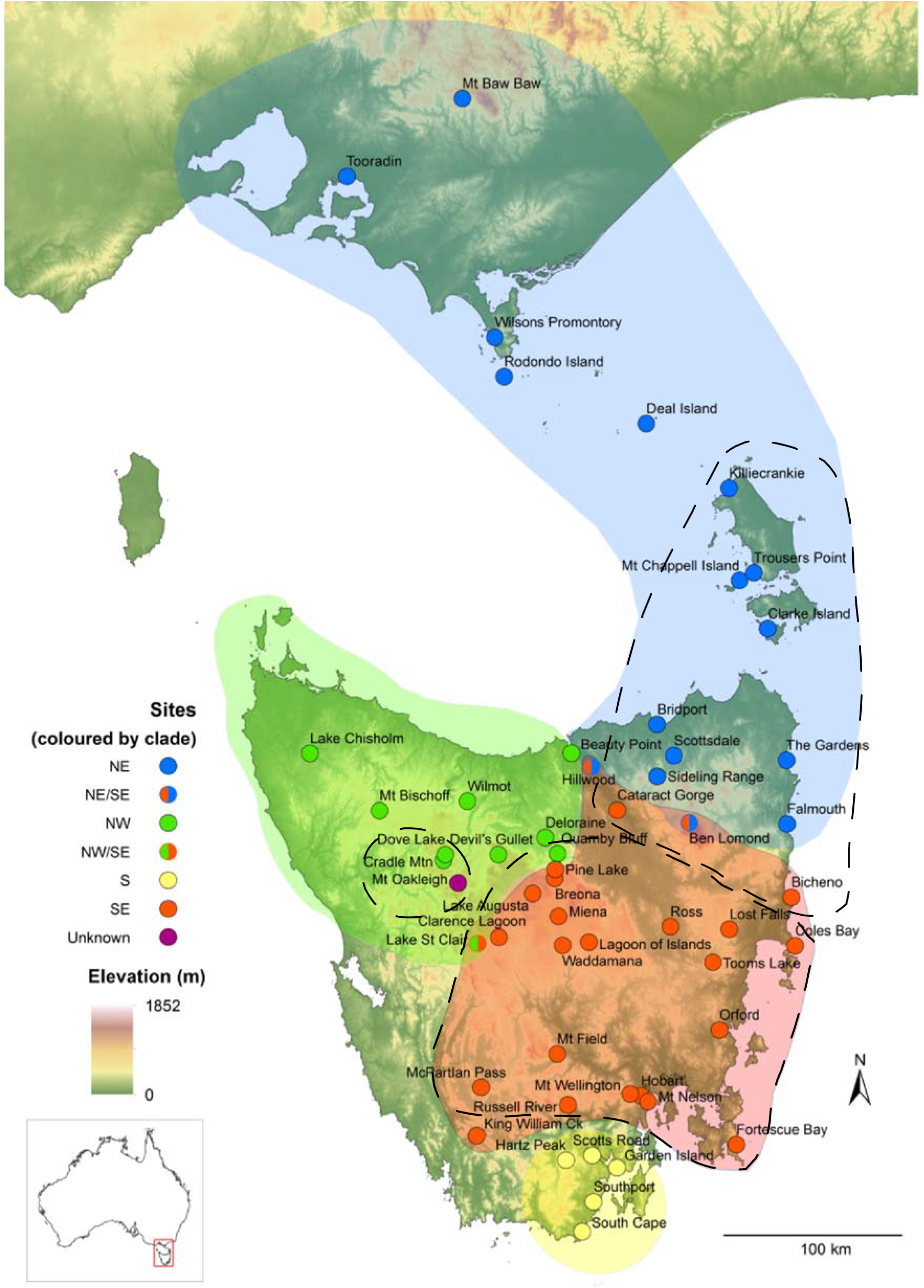
*Niveoscincus metallicus* sampling locations coloured by four major regional mitochondrial clades. Sites representing two clades are shown by circles with two colours. The geographic distribution of mitochondrially-defined clades in *N. ocellatus* is indicated by dashed lines. Base map: 250 m DEM (GeoScience Australia).

The divergence dating tree recovered the same clades as the non ultrametric tree (Supplementary Figure 1). The divergence of the northwest clade was placed at 10.0 Ma (95% HPD: 4.5–18.4 Ma). The southern clade diverged around 5.8 Ma (2.5–10.6 Ma), and the split between the northeast and southeast clades occurred around 4.4 Ma (1.9–8.1 Ma). Within the major clades, deep internal branches diverged approximately 2.8–1.5 Ma. Within the northeast clade, the divergence between Tasmania and the group comprising the Bass Strait islands and mainland Australia was dated to around 2.8 Ma (1.2–5.2 Ma), while mainland Australian populations diverged 1–2 Ma.

The greatest gain in mitochondrial Φ_CT_ occurred at K = 4 groups in SAMOVA, and the assignment of sampling sites to groups was equivalent to the four major mitochondrial clades resolved by phylogenetic analyses. High and significant levels of population structure were detected among all comparisons tested using AMOVA: among all sampling sites, among sites within regional clades, and among regional clades, with the highest proportion of variation (63%) attributable to regional clades (Table 2). Each regional clade was also significantly more structured than would be expected by chance alone, with the exception of the Northwestern clade, in which considerably more variation was contained within populations rather than among populations (Table 2). The Northeastern clade was the most highly structured clade, followed by the Southeastern and Southern clades (Table 2). AMOVA was also conducted on groups of populations within the Northeastern clade (Table 2); the island populations were highly structured, mainland Australian populations were somewhat less structured and north-eastern Tasmanian populations were significantly but comparatively weakly structured (Table 2).

**Table 2.** Analysis of molecular variance (AMOVA) comparing mitochondrial and nuclear genetic structure within sampling sites and among and between groups defined by major mitochondrial clades in *Niveoscincus metallicus*. Analyses were conducted with and without admixed sites (sites representing more than one mitochondrial lineage: Ben Lomond, Hillwood and Lake St Clair).

| Source of variation | mtDNA |  |  | nDNA |  |  |
| --- | --- | --- | --- | --- | --- | --- |
|  | Fixation index | % total | P-value | Fixation index | % total | P-value |
| **All samples** |  |  |  |  |  |  |
| – *with admixed sites* |  |  |  |  |  |  |
| Among regional groups | 0.63 ($\Phi_{CT}$) | 63.41 | <0.0001 | 0.15 ($\Phi_{CT}$) | 14.62 | 0.0230 |
| Among sites within regional groups | 0.62 ($\Phi_{SC}$) | 23.04 | <0.0001 | 0.65 ($\Phi_{SC}$) | 55.47 | <0.0001 |
| Among sites | 0.86 ($\Phi_{ST}$) | 13.55 | <0.0001 | 0.70 ($\Phi_{ST}$) | 29.91 | <0.0001 |
| – *without admixed sites* |  |  |  |  |  |  |
| Among regional groups | 0.61 ($\Phi_{CT}$) | 64.44 | <0.0001 | | | |
| Among sites within regional groups | 0.62 ($\Phi_{SC}$) | 22.04 | <0.0001 | | | |
| Among sites | 0.86 ($\Phi_{ST}$) | 13.52 | <0.0001 | | | |
| **Southeast clade** |  |  |  |  |  |  |
| – *with admixed sites* |  |  |  |  |  |  |
| Among populations | 0.64 ($\Phi_{ST}$) | 63.96 | <0.0001 | 0.64 ($\Phi_{ST}$) | 63.75 | <0.0001 |
| – *without admixed sites* |  |  |  |  |  |  |
| Among populations | 0.62 ($\Phi_{ST}$) | 61.68 | <0.0001 | | | |
| **Northwest clade** |  |  |  |  |  |  |
| – *with admixed sites* |  |  |  |  |  |  |
| Among populations | 0.15 ($\Phi_{ST}$) | 14.55 | 0.0639 | 0.43 ($\Phi_{ST}$) | 42.63 | <0.0001 |
| – *without admixed sites* |  |  |  |  |  |  |
| Among populations | 0.14 ($\Phi_{ST}$) | 14.04 | 0.0599 | | | |
| **Southern clade** |  |  |  |  |  |  |
| Among populations | 0.47 ($\Phi_{ST}$) | 47.94 | <0.0001 | 0.21 ($\Phi_{ST}$) | 21.06 | 0.0028 |
| **Northeast clade** |  |  |  |  |  |  |
| – *with admixed sites* |  |  |  |  |  |  |
| Among populations | 0.79 ($\Phi_{ST}$) | 79.03 | <0.0001 | 0.89 ($\Phi_{ST}$) | 89.44 | <0.0001 |
| – *without admixed sites* |  |  |  |  |  |  |
| Among populations | 0.79 ($\Phi_{ST}$) | 79.17 | <0.0001 | | | |

**Table 3.**
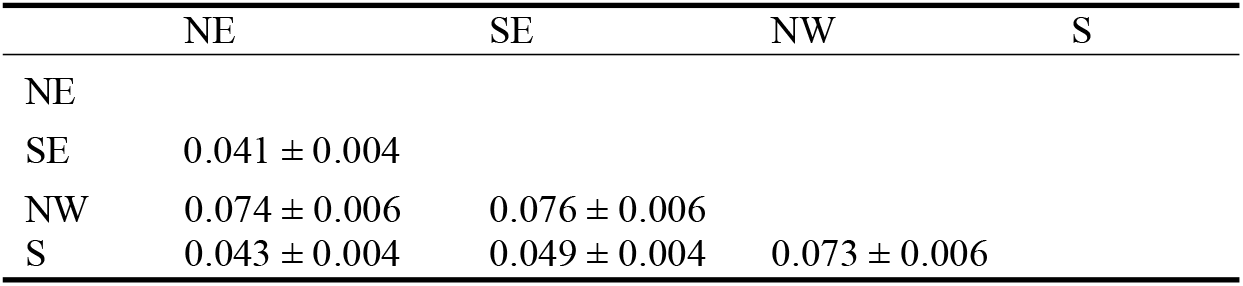
Uncorrected mean genetic distances (± standard error from 500 bootstrap replicates) between regional clades detected from phylogenetic analysis of 1406 base pairs of ND2 and ND4 mitochondrial sequence for *Niveoscincus metallicus*.

### Nuclear variation

Nuclear (β-globin) sequences were much less variable than mitochondrial sequences, with 87 polymorphic and 44 parsimony informative nucleotide sites from a total of 782 bp. In addition, there were three single nucleotide indels, and longer indels of 12, 13, 14 and 16 bp. A total of 83 unique β-globin haplotypes were recovered from 168 alleles (84 individuals) from 34 locations across the species range (Table 1). Several haplotypes represented multiple individuals from several sampling locations, and network analysis revealed little phylogeographic structure (Supplementary Figure 2). During SAMOVA the greatest increase in Φ_CT_ was detected at K = 2, with the mainland Australian sites plus Rodondo Island (north Bass Strait) and Waddamana (south Central Plateau) grouped together, and all remaining sites allocated to the other group. Significant levels of population structure were detected by AMOVA based on the mitochondrially-defined groups: among regional groups, among populations within groups, and among populations (Table 2). The relative amount of structuring of each major regional group was mostly consistent with mtDNA. The northeast clade had the highest level of population genetic structure, followed by the southeast clade. In contrast to the mitochondrial results, the northwest clade was significantly structured and was more highly structured than the southern clade (Table 2).

## Discussion

We revealed significant structuring of genetic variation across the range of *N. metallicus*, including distinct mtDNA clades corresponding to Tasmania’s northwest, northeast (including Bass Strait islands and mainland Australia), southeast, and south. The phylogenetic network recovered from nuclear sequence data failed to resolve the same groups, but revealed significant regional partitioning of variation consistent with the mtDNA pattern. The dated mtDNA phylogeny indicates that the *N. metallicus* regional groups were established prior to the Pleistocene, and hence persisted through multiple glacial-interglacial cycles. The broad mitochondrial phylogeographic patterns and corresponding divergence timescales are strikingly similar to those found in *N. ocellatus* (Figure 3; Cliff *et al.*, 2015). These species are closely related, have similar distributions and thermal niches (Caldwell *et al.*, 2017), and differ mainly in their dominant habitat, with *N. ocellatus* being obligate saxicolous (Wapstra *et al.*, 1999), while *N. metallicus* is typically found in close proximity to deep litter or other shelter such as fallen logs or rocks (Rawlinson, 1974; Melville & Swain, 1999). Therefore, the concordant mtDNA patterns in these species are more likely to reflect a shared underlying mechanism (phylogeographic parallelism) than chance coincidence (phylogeographic convergence).

Isolation into multiple Pleistocene refugia is the most parsimonious explanation for the presence of regional phylogeographic structuring within Tasmanian *Niveoscincus*. This hypothesis is supported by the presence of contact zones between divergent lineages at high elevation areas that experienced glacial and periglacial activity during the LGM (Colhoun *et al.*, 1996; Taberlet *et al.*, 1998; Barrows *et al.*, 2002; Hewitt, 2011). Similarly, divergence between haplotypes in the high elevation eastern Central Plateau was shallow, suggesting recent colonisation and expansion into this region. During Pleistocene glaciations the distribution of *N. metallicus* was probably limited by cold conditions and the absence of suitable habitat across large areas of its current range, isolating populations into regional refugia. The effect of temperature on reproduction in widespread lowland and restricted alpine *Niveoscincus* species in Tasmania is well studied (Jones & Swain, 1996; Swain & Jones, 2000b; Swain & Jones, 2000a; Atkins *et al.*, 2007; Wapstra *et al.*, 2009; Cadby *et al.*, 2010; Pen *et al.*, 2010; Uller *et al.*, 2011), and a mechanistic link between thermally limited reproduction and distribution has been established (Olsson & Shine, 1998, 1999; Wapstra *et al.*, 1999). The distribution of generalist lowland *Niveoscincus* (*N. ocellatus* and *N. metallicus*) is currently limited by cold conditions in high elevation (~1100-1200 m) areas (Wapstra *et al.*, 1999), and cold, alpine environments were more extensive during Pleistocene glaciations (Kirkpatrick & Fowler, 1998; Colhoun & Barrows, 2011; Colhoun & Shimeld, 2012). LGM temperature depressions were up to 6.5°C (Colhoun, 1985; Fletcher & Thomas, 2010) and tree-lines were close to the modern day sea level in places (Gibson *et al.*, 1987).

The absence or contraction of vegetation communities during Pleistocene climate cycles may have also affected the distribution of *N. metallicus*. This species does not occur in closed wet forest and is absent above the subalpine tree-line (Melville & Swain, 1999). During Pleistocene glaciations, the dominant woodland and open forest habitat of *N. metallicus* was displaced by semiarid grassland and extensive areas of subalpine vegetation in Tasmania and southern Australia (Kirkpatrick & Fowler, 1998; Kershaw *et al.*, 2007; Colhoun & Shimeld, 2012). However, vegetation changes were unlikely to influence the distribution of rock habitat for *N. ocellatus*, and given concordant phylogeographic patterns with *N. metallicus*, it would appear that climate was the main driver of past distributions in both species.

The divergences between major *N. metallicus* clades appear to predate the LGM, and have been maintained across multiple favourable interglacial periods when population connections may have been established. Glacial periods may have reinforced or resurrected isolation of these lineages following interglacial periods. For instance, our present day (interglacial) sampling revealed peripheral overlap of the major *N. metallicus* clades, and this has also been observed for *N. ocellatus* following fine-scale sampling along a transect across clade boundaries (Burridge, unpublished). Similarly, the major phylogeographic breaks do not reflect contemporary landscape features that would restrict dispersal in either of these species, which are patchily but more-or-less continuously distributed with the exception of areas above 1100 m elevation. However, the most likely areas for lineage exchange correspond to central and high-elevation areas of Tasmania that would have been most affected by Pleistocene glaciation. Lineages exchanged at their spatial peripheries during previous interglacials may have failed to be retained in subsequent low elevation glacial refugia, hence reinforcing prior phylogeographic structuring. High-density blocking may have also reduced the invasion of lineages from adjacent clades during interglacials (Hewitt, 2011; Waters, 2011; Waters *et al.*, 2013). Intra-specific competition can also maintain phylogeographic structuring. For example, victory in aggressive encounters between adult male *N. microlepidotus* is strongly correlated with territorial residency, with 72% of conflicts resulting in the losing male leaving the contested territory (Olsson & Shine, 2000). However, competition among females is required with respect to mtDNA structuring.

In northwest Tasmania there is strong evidence for the long-term isolation of lineages across a diverse range of taxa, with candidate isolating mechanisms already identified. Distinct lineages are observed for *N. ocellatus* (Cliff *et al.*, 2015), Tasmanian pademelon (Macqueen *et al.*, 2009), Tasmanian devil (Jones *et al.*, 2004; Brüniche-Olsen *et al.*, 2014), Eastern quoll (Cardoso *et al.*, 2014), Tasmanian tree frog (Zhang *et al.*, 2014), White’s skink (Chapple *et al.*, 2005) and the canopy tree species *Eucalyptus obliqua* (Bloomfield *et al.*, 2011). The isolation of these lineages may have been facilitated or reinforced by glaciers in the Mersey and Forth River valleys that extended to within 20 km of the present-day coastline (Colhoun *et al.*, 1996). During glacial periods, the low lying central north of Tasmania was probably also arid and treeless (Kirkpatrick & Fowler, 1998), further inhibiting movement in a range of taxa, although pollen records are lacking for this region (Colhoun & Shimeld, 2012). In contrast, Kirkpatrick and Fowler (1998) suggested that northwest Tasmania was a refugium for sclerophyll woodland at the LGM, which is corroborated by pollen evidence (Colhoun & Shimeld, 2012), and would promote distinct northwest lineages of those taxa listed above.

Genetic isolation between the remaining major clades may also reflect utilisation of distinct refugia during Pleistocene glaciations. In particular, accumulating evidence suggests that northeast Tasmania was an important glacial refugium for a number of species. The region supports genetically distinct populations within the rainforest tree species *Nothofagus cunninghamii* (Worth *et al.*, 2009; Sheedy *et al.*, 2015) and *Atherosperma moschatum* (Worth *et al.*, 2011), and several eucalypt species (McKinnon *et al.*, 2004; Nevill *et al.*, 2010; Gauli *et al.*, 2014; Nevill *et al.*, 2014). Genetic evidence and vegetation modelling indicate that the eastern foothills of the Ben Lomond Plateau supported eucalypt woodland at the LGM (Kirkpatrick & Fowler, 1998; Gauli *et al.*, 2014). However, immediately to the south, LGM pollen records indicate extensive areas of semi-arid grassland extending inland from the central east coast (Colhoun & Shimeld, 2012; Mackenzie & Moss, 2014). The Midlands region was probably also more arid and treeless during glaciations (Sigleo & Colhoun, 1981; Kirkpatrick & Fowler, 1998; Colhoun & Shimeld, 2012), and may have been unsuitable for persistence of *N. metallicus*, but a stronger understanding of its physiological responses and tolerance of *N. metallicus* to aridity is required to infer potential responses to aridity-driven environmental changes (Caldwell *et al.*, 2015). Refugia would have also existed to maintain the Southeast and South clades. The presence of non-alpine adapted *Niveoscincus* on Pedra Branca, an islet on the continental shelf to the south of Tasmania, indicates persistence during the time of glacial low sea stands (Rounsevell *et al.*, 1985).

Despite concordance in the broad regional phylogeographic patterns in *N. metallicus* and *N. ocellatus*, a striking difference occurred within northeast Tasmania. In contrast to *N. ocellatus*, there was a lack of lowland site monophyly for mtDNA sequence variation within the Northeastern clade for *N. metallicus*. This may reflect differences in habitat requirements in areas continuously occupied during the Pleistocene. As *N. metallicus* is more a habitat generalist, it would therefore experience greater effective population sizes and gene flow, and consequently less phylogeographic structuring. Similar explanations have been offered for differences in gene flow and population structure in other co-distributed reptile species (Haines *et al.*, 2014; Podnar *et al.*, 2014)

In Tasmanian *N. metallicus*, phylogeographic structuring has been maintained when populations were potentially connected during interglacials. Populations from the Bass Strait islands and mainland Australia were also individually divergent and monophyletic, but these regions were isolated during interglacials and terrestrially connected during glacial periods (marine transgressions). For example, during the last glacial period, this land bridge was exposed from about 42 ka to 14 ka (Lambeck & Chappell, 2001). Similar to the explanation for phylogeographic structuring in Tasmania, mixing between Bass Strait island and mainland Australian lineages may have been inhibited during low sea stands by high density blocking or intraspecific competition, and signatures of any mixing may have been lost during (interglacial) isolation caused by elevated sea levels. Alternatively, the Bassian Isthmus may have represented a formidable barrier to dispersal for this species. Aridity during late Pleistocene glacial cycles (Kershaw *et al.*, 2003; McLaren & Wallace, 2010; McLaren *et al.*, 2012) may have prevented the establishment of suitable habitat, restricting dispersal across the isthmus. Although evidence is limited, most of the Bassian Isthmus was probably semi-arid and dominated by grasslands at the LGM (D&Costa *et al.*, 1993; Kirkpatrick & Fowler, 1998). Either or both of these factors must have been significant for *Niveoscincus*, because phylogeographic studies of other reptiles across Bass Strait did not recover Tasmanian and Bass Strait populations as monophyletic, either with respect to one another (Fairbairn *et al.*, 1998; Dubey & Shine, 2010; Ng *et al.*, 2014), or relative to mainland Australian populations (Chapple *et al.*, 2005; Keogh *et al.*, 2005; Chapple *et al.*, 2011b). A faster rate of lineage sorting is unlikely to explain the greater phylogeographic structuring in these two *Niveoscincus* across Bass Strait, because their effective population sizes are unlikely to be substantially smaller than any of these other reptiles.

The presence of broad phylogeographic structuring within *N. ocellatus* and *N. metallicus* that predates the LGM is consistent with multiple studies conducted on amphibians and squamates distributed in southeast Australia (Byrne, 2008; Symula *et al.*, 2008; Dubey & Shine, 2010; Byrne *et al.*, 2011; Chapple *et al.*, 2011a; Chapple *et al.*, 2011b; Haines *et al.*, 2014; Ng *et al.*, 2014), and the temperate Southern Hemisphere more broadly (Wallis & Trewick, 2009; Sersic *et al.*, 2011; Breitman *et al.*, 2012; Vera-Escalona *et al.*, 2012; Turchetto-Zolet *et al.*, 2013). High haplotype and nucleotide diversity within all regional clades also suggests that *N. metallicus* populations persisted in multiple locations within each region, rather than being constricted to a single refugia from which they subsequently expanded. Hence, the emerging Southern Hemisphere pattern is one of persistence and diversification in multiple glacial refugia broadly across the current species range (Byrne, 2008; Byrne *et al.*, 2011; Haines *et al.*, 2014).

Southern Hemisphere patterns contrast with those reflecting widespread extirpation and low latitude refugia, as observed for several Northern Hemisphere taxa (Taberlet *et al.*, 1998; but see Stewart & Lister, 2001; Hewitt, 2004; Stewart *et al.*, 2010; Hewitt, 2011). However, recent work in southern regions of Europe and North America has also revealed increasingly complex phylogeographic patterns indicative of long-term persistence in multiple refugia (Canestrelli *et al.*, 2007; Gómez & Lunt, 2007; Nieto Feliner, 2011; Canestrelli *et al.*, 2012; Podnar *et al.*, 2014; Bellati *et al.*, 2015). It appears that the paradigm of high latitude extirpation and long-range postglacial re-colonisation from a small number of low latitude refugia does not apply universally, and that complex, localised and species-specific responses to Pleistocene climatic oscillations are the general expectation in unglaciated and comparatively lightly glaciated temperate regions (Soltis *et al.*, 2006; Byrne, 2008; Wallis & Trewick, 2009; Byrne *et al.*, 2011; Turchetto-Zolet *et al.*, 2013). However, here we also show that major phylogeographic breaks can be shared by similar species, suggesting the operation of a common mechanism.

## ACKNOWLEDGEMENTS

We are grateful to staff at the CSIRO Australian National Wildlife Collection, the South Australian Museum and Victoria Museum, as well as C. Jennings (Australian National University) and M. Driessen (DPIPWE Tasmania), for assistance in accessing some of the samples used in this study. We would like to thank the volunteers who helped collect samples in the field: M. Caldwell, A. Demir, T. Feldmanis, J. Gruber, K. Munch, P. Richardson and E. K. Yuni, with particular thanks to R. De Paoli. We would also like to thank T. Wenner and A. Pracejus for valuable assistance in the laboratory, J. Kirkpatrick, G. Jordon, J. Worth for insights on biotic responses to glaciations in Tasmania, and C. Moritz and D. Rosauer for technical support in the production of the map included in this publication. Samples were collected with approval from the University of Tasmania Animal Ethics Committee (permit A12286). This research was funded by the Australian Research Council (FT110100597).

## References

Araújo, M.B., Nogués-Bravo, D., Diniz-Filho, J.A.F., Haywood, A.M., Valdes, P.J. & Rahbek, C. (2008) Quaternary climate changes explain diversity among reptiles and amphibians. Ecography, 31, 8–15.

Atkins, N., Swain, R.O.Y., Wapstra, E. & Jones, S.M. (2007) Late stage deferral of parturition in the viviparous lizard *Niveoscincus ocellatus* (Gray 1845): implications for offspring quality and survival. Biological Journal of the Linnean Society, 90, 735–746.

Avise, J.C. (1992) Molecular population structure and the biogeographic history of a regional fauna-a case-history with lessons for conservation biology. Oikos, 63, 62–76.

Barrows, T.T., Stone, J.O., Fifield, L.K. & Cresswell, R.G. (2002) The timing of the Last Glacial Maximum in Australia. Quaternary Science Reviews, 21, 159–173.

Beheregaray, L.B. (2008) Twenty years of phylogeography: the state of the field and the challenges for the Southern Hemisphere. Molecular Ecology, 17, 3754–3774.

Bell, R.C., Parra, J.L., Tonione, M., Hoskin, C.J., Mackenzie, J.B., Williams, S.E. & Moritz, C. (2010) Patterns of persistence and isolation indicate resilience to climate change in montane rainforest lizards. Molecular Ecology, 19, 2531–2544.

Bellati, A., Carranza, S., Garcia-Porta, J., Fasola, M. & Sindaco, R. (2015) Cryptic diversity within the Anatololacerta species complex (Squamata: Lacertidae) in the Anatolian Peninsula: Evidence from a multi-locus approach. Molecular Phylogenetics and Evolution, 82, 219–233.

Bensasson, D., Zhang, D.-X., Hartl, D.L. & Hewitt, G.M. (2001) Mitochondrial pseudogenes: evolution’s misplaced witnesses. Trends in Ecology & Evolution, 16, 314–321.

Bermingham, E. & Avise, J.C. (1986) Molecular zoogeography of freshwater fishes in the southeastern United States. Genetics, 113, 939–965.

Blois, J.L., McGuire, J.L. & Hadly, E.A. (2010) Small mammal diversity loss in response to late-Pleistocene climatic change. Nature, 465, 771–775.

Bloomfield, J.A., Nevill, P., Potts, B.M., Vaillancourt, R.E. & Steane, D.A. (2011) Molecular genetic variation in a widespread forest tree species Eucalyptus obliqua (Myrtaceae) on the island of Tasmania. Australian Journal of Botany, 59, 226–237.

Bouckaert, R., Heled, J., Kuhnert, D., Vaughan, T., Wu, C.H., Xie, D., Suchard, M.A., Rambaut, A. & Drummond, A.J. (2014) BEAST2: A Software Platform for Bayesian Evolutionary Analysis. PLoS Computational Biology, 10

Breitman, M.F., Avila, L.J., Sites, J.W. & Morando, M. (2012) How lizards survived blizzards: phylogeography of the Liolaemus lineomaculatus group (Liolaemidae) reveals multiple breaks and refugia in southern Patagonia and their concordance with other codistributed taxa. Molecular Ecology, 21, 6068–6085.

Brüniche-Olsen, A., Jones, M.E., Austin, J.J., Burridge, C.P. & Holland, B.R. (2014) Extensive population decline in the Tasmanian devil predates European settlement and devil facial tumour disease.

Burridge, C.P., Craw, D., Jack, D.C., King, T.M. & Waters, J.M. (2008) Does fish ecology predict dispersal across a river drainage divide? Evolution, 62, 1484–1499.

Byrne, M. (2008) Evidence for multiple refugia at different time scales during Pleistocene climatic oscillations in southern Australia inferred from phylogeography. Quaternary Science Reviews, 27, 2576–2585.

Byrne, M., Yeates, D.K., Joseph, L., Kearney, M., Bowler, J., Williams, M.A.J., Cooper, S., Donnellan, S.C., Keogh, J.S., Leys, R., Melville, J., Murphy, D.J., Porch, N. & Wyrwoll, K.H. (2008) Birth of a biome: insights into the assembly and maintenance of the Australian arid zone biota. Molecular Ecology, 17, 4398–4417.

Byrne, M., Steane, D.A., Joseph, L., Yeates, D.K., Jordan, G.J., Crayn, D., Aplin, K., Cantrill, D. J., Cook, L.G., Crisp, M.D., Keogh, J.S., Melville, J., Moritz, C., Porch, N., Sniderman, J.M.K., Sunnucks, P. & Weston, P.H. (2011) Decline of a biome: evolution, contraction, fragmentation, extinction and invasion of the Australian mesic zone biota. Journal of Biogeography, 38, 1635–1656.

Cadby, C.D., While, G.M., Hobday, A.J., Uller, T. & Wapstra, E. (2010) Multi-scale approach to understanding climate effects on offspring size at birth and date of birth in a reptile. Integrative Zoology, 5, 164–175.

Caldwell, A.J., While, G.M. & Wapstra, E. (2017) Plasticity of thermoregulatory behaviour in response to the thermal environment by widespread and alpine reptile species. Animal Behaviour, 132, 217–227.

Caldwell, A.J., While, G.M., Beeton, N.J. & Wapstra, E. (2015) Potential for thermal tolerance to mediate climate change effects on three members of a cool temperate lizard genus, Niveoscincus. Journal of Thermal Biology, 52, 14–23.

Camargo, A., Sinervo, B. & Sites Jr, J.W. (2010) Lizards as model organisms for linking phylogeographic and speciation studies. Molecular Ecology, 19, 3250–3270.

Canestrelli, D., Cimmaruta, R. & Nascetti, G. (2007) Phylogeography and historical demography of the Italian treefrog, Hyla intermedia, reveals multiple refugia, population expansions and secondary contacts within peninsular Italy. Molecular Ecology, 16, 4808–4821.

Canestrelli, D., Salvi, D., Maura, M., Bologna, M.A. & Nascetti, G. (2012) One Species, Three Pleistocene Evolutionary Histories: Phylogeography the Italian Crested Newt, Triturus carnifex. PLoS one, 7

Cardoso, M.J., Mooney, N., Eldridge, M.D.B., Firestone, K.B. & Sherwin, W.B. (2014) Genetic monitoring reveals significant population structure in eastern quolls: implications for the conservation of a threatened carnivorous marsupial. Australian Mammalogy, 36, 169–177.

Carstens, B.C., Brunsfeld, S.J., Demboski, J.R., Good, J.M. & Sullivan, J. (2005) Investigating the evolutionary history of the Pacific Northwest mesic forest ecosystem: hypothesis testing within a comparative phylogeographic framework. Evolution, 59, 1639–1652.

Chapple, D.G., Keogh, J.S. & Hutchinson, M.N. (2005) Substantial genetic substructuring in southeastern and alpine Australia revealed by molecular phylogeography of the *Egernia whitii* (Lacertilia: Scincidae) species group. Molecular Ecology, 14, 1279–1292.

Chapple, D.G., Chapple, S.N.J. & Thompson, M.B. (2011a) Biogeographic barriers in south-eastern Australia drive phylogeographic divergence in the garden skink, Lampropholis guichenoti. Journal of Biogeography, 38, 1761–1775.

Chapple, D.G., Hoskin, C.J., Chapple, S.N.J. & Thompson, M.B. (2011b) Phylogeographic divergence in the widespread delicate skink (*Lampropholis delicata*) corresponds to dry habitat barriers in eastern Australia. BMC Evolutionary Biology, 11, 191–208.

Clement, M., Posada, D. & Crandall, K.A. (2000) TCS: a computer program to estimate gene genealogies. Molecular Ecology, 9, 1657–1659.

Cliff, H., Burridge, C.P. & Wapstra, E. (2015) Persistence and dispersal in a Southern Hemiphere glaciated landscape: the phylogeography of the spotted snow skink (*Niveoscincus ocellatus*) in Tasmania. BMC Evolutionary Biology, in press

Colhoun, E. & Barrows, T. (2011) The Glaciation of Australia. Quaternary Glaciations-Extent and Chronology: A closer look (ed. by J. Ehlers, P.L. Gibbard and P.D. Hughes), pp. 1037–1045. Elsevier, Amsterdam, Netherlands.

Colhoun, E. & Shimeld, P. (2012) Late-Quaternary vegetation history of Tasmania from pollen records. Peopled Landscapes: archaeological and biogeographical approaches to landscapes (ed. by S. Haberle and B. David). ANU Press, Canberra.

Colhoun, E.A. (1985) Pre-Last Glacial Maximum vegetation history at Henty Bridge, Western Tasmania. New Phytologist, 100, 681–690.

Colhoun, E.A., Hannan, D. & Kiernan, K. (1996) Late Wisconsin glaciation of Tasmania. Papers and Proceedings of the Royal Society of Tasmania (ed by, pp. 33–45

D’Costa, D.M., Grindrod, J. & Ogden, R. (1993) Preliminary environmental reconstructions from late Quaternary pollen and mollusc assemblages at Egg Lagoon, King Island, Bass Strait. Australian Journal of Ecology, 18, 351–366.

Darriba, D., Taboada, G.L., Doallo, R. & Posada, D. (2012) jModelTest 2: more models, new heuristics and parallel computing. Nature Methods, 9, 772–772.

Dawson, M.N. (2012) Parallel phylogeographic structure in ecologically similar sympatric sister taxa. Molecular Ecology, 21, 987–1004.

Dawson, M.N., Hays, C.G., Grosberg, R.K. & Raimondi, P.T. (2014) Dispersal potential and population genetic structure in the marine intertidal of the eastern North Pacific. Ecological Monographs, 84, 435–456.

Dawson, M.N., Louie, K.D., Barlow, M., Jacobs, D.K. & Swift, C.C. (2002) Comparative phylogeography of sympatric sister species, *Clevelandia ios* and *Eucyclogobius newberryi* (Teleostei, Gobiidae), across the California Transition Zone. Molecular Ecology, 11, 1065–1075.

de Lafontaine, G., Amasifuen Guerra, C.A., Ducousso, A. & Petit, R.J. (2014) Cryptic no more: soil macrofossils uncover Pleistocene forest microrefugia within a periglacial desert. New Phytologist, 204, 715–729.

DeChaine, E.G. & Martin, A.P. (2006) Using coalescent simulations to test the impact of quaternary climate cycles on divergence in an alpine plant-insect association. Evolution, 60, 1004–1013.

Drummond, A.J. & Suchard, M.A. (2010) Bayesian random local clocks, or one rate to rule them all. Bmc Biology, 8

Dubey, S. & Shine, R. (2010) Evolutionary Diversification of the Lizard Genus Bassiana (Scincidae) across Southern Australia. PLoS one, 5

Dupanloup, I., Schneider, S. & Excoffier, L. (2002) A simulated annealing approach to define the genetic structure of populations. Molecular Ecology, 11, 2571–2581.

Excoffier, L. & Lischer, H.E.L. (2010) Arlequin suite ver 3.5: a new series of programs to perform population genetics analyses under Linux and Windows. Molecular Ecology Resources, 10, 564–567.

Excoffier, L., Smouse, P.E. & Quattro, J.M. (1992) Analysis of Molecular Variance Inferred from Metric Distances among DNA Haplotypes-Application to Human Mitochondrial-DNA Restriction Data. Genetics, 131, 479–491.

Fairbairn, J., Shine, R., Moritz, C. & Frommer, M. (1998) Phylogenetic Relationships between Oviparous and Viviparous Populations of an Australian Lizard (Lerista bougainvillii, Scincidae). Molecular Phylogenetics and Evolution, 10, 95–103.

Fletcher, M.-S. & Thomas, I. (2010) A quantitative Late Quaternary temperature reconstruction from western Tasmania, Australia. Quaternary Science Reviews, 29, 2351–2361.

Flot, J.F. (2010) SEQPHASE: a web tool for interconverting phase input/output files and fasta sequence alignments. Molecular Ecology Resources, 10, 162–166.

Fordham, D.A., Brook, B.W., Moritz, C. & Nogués-Bravo, D. (2014) Better forecasts of range dynamics using genetic data. Trends in Ecology & Evolution, 29, 436–443.

Gauli, A., Steane, D.A., Vaillancourt, R.E. & Potts, B.M. (2014) Molecular genetic diversity and population structure in Eucalyptus pauciflora subsp pauciflora (Myrtaceae) on the island of Tasmania. Australian Journal of Botany, 62, 175–188.

Gibson, N., Kiernan, K. & Macphail, M. (1987) A fossil bolster plant from the King River, Tasmania. Papers and Proceedings of the Royal Society of Tasmania, 121, 35–42.

Gómez, A. & Lunt, D. (2007) Refugia within Refugia: Patterns of Phylogeographic Concordance in the Iberian Peninsula. Phylogeography of Southern European Refugia (ed. by S. Weiss and N. Ferrand), pp. 155–188. Springer Netherlands.

Guindon, S. & Gascuel, O. (2003) A simple, fast, and accurate algorithm to estimate large phylogenies by maximum likelihood. Systematic Biology, 52, 696–704.

Haines, M.L., Moussalli, A., Stuart-Fox, D., Clemann, N. & Melville, J. (2014) Phylogenetic evidence of historic mitochondrial introgression and cryptic diversity in, the genus Pseudemoia (Squamata: Scincidae). Molecular Phylogenetics and Evolution, 81, 86–95.

Hare, K.M., Daugherty, C.H. & Chapple, D.G. (2008) Comparative phylogeography of three skink species (*Oligosoma moco, O. smithi, O. suteri*; Reptilia: Scincidae) in northeastern New Zealand. Molecular Phylogenetics and Evolution, 46, 303–315.

Hewitt, G.M. (2000) The genetic legacy of the Quaternary ice ages. Nature, 405, 907–913.

Hewitt, G.M. (2004) Genetic consequences of climatic oscillations in the Quaternary. Philosophical Transactions of the Royal Society of London Series B-Biological Sciences, 359, 183–195.

Hewitt, G.M. (2011) Quaternary phylogeography: the roots of hybrid zones. Genetica, 139, 617–638.

Hickerson, M.J. & Cunningham, C.W. (2005) Contrasting Quaternary histories in an ecologically divergent sister pair of low-dispersing intertidal fish (*Xiphister*) revealed by multilocus DNA analysis. Evolution, 59, 344–360.

Huson, D.H. & Bryant, D. (2006) Application of phylogenetic networks in evolutionary studies. Molecular Biology and Evolution, 23, 254–267.

Jackson, S.T. & Overpeck, J.T. (2000) Responses of plant populations and communities to environmental changes of the late Quaternary. Paleobiology, 26, 194–220.

Jones, M.E., Paetkau, D., Geffen, E.L.I. & Moritz, C. (2004) Genetic diversity and population structure of Tasmanian devils, the largest marsupial carnivore. Molecular Ecology, 13, 2197–2209.

Jones, S.M. & Swain, R. (1996) Annual reproductive cycle and annual cycles of reproductive hormones in plasma of female *Niveoscincus metallicus* (Scincidae) from Tasmania. Journal of Herpetology, 30, 140–146.

Keogh, J.S., Scott, I.A.W. & Hayes, C. (2005) Rapid and repeated origin of insular gigantism and dwarfism in Australian tiger snakes. Evolution, 59, 226–233.

Kershaw, A.P., McKenzie, G.M., Porch, N., Roberts, R.G., Brown, J., Heijnis, H., Orr, M.L., Jacobsen, G. & Newallt, P.R. (2007) A high-resolution record of vegetation and climate through the last glacial cycle from Caledonia Fen, southeastern highlands of Australia. Journal of Quaternary Science, 22, 481–500.

Kershaw, P., Moss, P. & Van Der Kaars, S. (2003) Causes and consequences of long-term climatic variability on the Australian continent. Freshwater Biology, 48, 1274–1283.

Kirkpatrick, J.B. & Fowler, M. (1998) Locating likely glacial forest refugia in Tasmania using palynological and ecological information to test alternative climatic models. Biological Conservation, 85, 171–182.

Lambeck, K. & Chappell, J. (2001) Sea Level Change Through the Last Glacial Cycle. Science, 292, 679–686.

Leigh, J.W. & Bryant, D. (2015) PopART: Full-feature software for haplotype network construction. Methods in Ecology and Evolution, 6, 1110–1116.

Lemmon, A.R., Brown, J.M., Stanger-Hall, K. & Lemmon, E.M. (2009) The Effect of Ambiguous Data on Phylogenetic Estimates Obtained by Maximum Likelihood and Bayesian Inference. Systematic Biology, 58, 130–145.

Lessa, E.P., Cook, J.A. & Patton, J.L. (2003) Genetic footprints of demographic expansion in North America, but not Amazonia, during the Late Quaternary. Proceedings of the National Academy of Sciences of the United States of America, 100, 10331–10334.

Lorenzen, E.D., Heller, R. & Siegismund, H.R. (2012) Comparative phylogeography of African savannah ungulates. Molecular Ecology, 21, 3656–3670.

Mackenzie, L. & Moss, P. (2014) A late Quaternary record of vegetation and climate change from Hazards Lagoon, eastern Tasmania. Quaternary International,

Macqueen, P., Goldizen, A.W. & Seddon, J.M. (2009) Response of a southern temperate marsupial, the Tasmanian pademelon (*Thylogale billardierii*), to historical and contemporary forest fragmentation. Molecular Ecology, 18, 3291–3306.

McKinnon, G.E., Jordan, G.J., Vaillancourt, R.E., Steane, D.A. & Potts, B.M. (2004) Glacial refugia and reticulate evolution: the case of the Tasmanian eucalypts. Philosophical Transactions of the Royal Society of London. Series B: Biological Sciences, 359, 275–284.

McLaren, S. & Wallace, M.W. (2010) Plio-Pleistocene climate change and the onset of aridity in southeastern Australia. Global and Planetary Change, 71, 55–72.

McLaren, S., Wallace, M.W. & Reynolds, T. (2012) The Late Pleistocene evolution of palaeo megalake Bungunnia, southeastern Australia: A sedimentary record of fluctuating lake dynamics, climate change and the formation of the modern Murray River. Palaeogeography Palaeoclimatology Palaeoecology, 317, 114–127.

Melville, J. & Swain, R. (1997) Spatial separation in two sympatric skinks, *Niveoscincus microlepidotus* and *N. metallicus*, from Tasmania. Herpetologica, 53, 126–132.

Melville, J. & Swain, R. (1999) Habitat associations and natural history of the Tasmanian “snow skinks” (*Niveoscincus* spp.) Papers and Proceedings of the Royal Society ofTasmania, 133, 57–64.

Melville, J. & Swain, R. (2003) Evolutionary correlations between escape behaviour and performance ability in eight species of snow skinks (*Niveoscincus*: Lygosominae) from Tasmania. Journal of Zoology, 261, 79–89.

Metts, B.S., Winne, C.T., Scott, D.E., Gibbons, J.W., Greene, J.L., Buhlmann, K.A., Poppy, S., Mills, T., Tuberville, T.D., Ryan, T.J. & Leiden, Y. (2000) The Global Decline of Reptiles, Déjà Vu Amphibians: Reptile species are declining on a global scale. Six significant threats to reptile populations are habitat loss and degradation, introduced invasive species, environmental pollution, disease, unsustainable use, and global climate change. BioScience, 50, 653–666.

Muller, K. (2006) Incorporating information from length-mutational events into phylogenetic analysis. Molecular Phylogenetics and Evolution, 38, 667–676.

Nelson-Tunley, M., Morgan-Richards, M. & Trewick, S.A. (2016) Genetic diversity and gene flow in a rare New Zealand skink despite fragmented habitat in a volcanic landscape. Biological Journal of the Linnean Society, 119, 37–51.

Nevill, P., Després, T., Bayly, M., Bossinger, G. & Ades, P. (2014) Shared phylogeographic patterns and widespread chloroplast haplotype sharing in Eucalyptus species with different ecological tolerances. Tree Genetics & Genomes, 10, 1079–1092.

Nevill, P.G., Bossinger, G. & Ades, P.K. (2010) Phylogeography of the world’s tallest angiosperm, *Eucalyptus regnans*: evidence for multiple isolated Quaternary refugia. Journal of Biogeography, 37, 179–192.

Ng, J., Clemann, N., Chapple, S.J. & Melville, J. (2014) Phylogeographic evidence links the threatened ‘Grampians’ Mountain Dragon (Rankinia diemensis Grampians) with Tasmanian populations: conservation implications in south-eastern Australia. Conservation Genetics, 15, 363–373.

Nieto Feliner, G. (2011) Southern European glacial refugia: A tale of tales. Taxon, 60, 365–372.

Olsson, M. & Shine, R. (1998) Timing of parturition as a maternal care tactic in an Alpine lizard species. Evolution, 52, 1861–1864.

Olsson, M. & Shine, R. (1999) Plasticity in frequency of reproduction in an alpine lizard, *Niveoscincus microlepidotus*. Copeia, 794–796.

Olsson, M. & Shine, R. (2000) Ownership influences the outcome of male-male contests in the scincid lizard, *Niveoscincus microlepidotus*. Behavioral Ecology, 11, 587–590.

Papadopoulou, A. & Knowles, L.L. (2016) Toward a paradigm shift in comparative phylogeography driven by trait-based hypotheses. Proc Natl Acad Sci U S A, 113, 8018–24.

Pen, I., Uller, T., Feldmeyer, B., Harts, A., While, G.M. & Wapstra, E. (2010) Climate-driven population divergence in sex-determining systems. Nature, 468, 436–438.

Podnar, M., Bruvo Mađarić, B. & Mayer, W. (2014) Non-concordant phylogeographical patterns of three widely codistributed endemic Western Balkans lacertid lizards (Reptilia, Lacertidae) shaped by specific habitat requirements and different responses to Pleistocene climatic oscillations. Journal of Zoological Systematics and Evolutionary Research, 52, 119–129.

Qiu, Y.X., Fu, C.X. & Comes, H.P. (2011) Plant molecular phylogeography in China and adjacent regions: Tracing the genetic imprints of Quaternary climate and environmental change in the world’s most diverse temperate flora. Molecular Phylogenetics and Evolution, 59, 225–244.

Rambaut, A. (2014) FigTree v1.4.2.

Rambaut, A. & Drummond, A.J. (2013) Tracer v1.6.

Rawlinson, P.A. (1974) Biogeography and ecology of the reptiles of Tasmania and the Bass Strait area. Biogeography and Ecology of Tasmania (ed. by W. Williams), pp. 291–338, W. Junk, The Hague, Netherlands.

Ronquist, F., Teslenko, M., van der Mark, P., Ayres, D.L., Darling, A., Hohna, S., Larget, B., Liu, L., Suchard, M.A. & Huelsenbeck, J.P. (2012) MrBayes 3.2: Efficient Bayesian Phylogenetic Inference and Model Choice Across a Large Model Space. Systematic Biology, 61, 539–542.

Rounsevell, D., Brothers, N. & Holdsworth, N. (1985) The status and ecology of the Pedra Branca skink, *Pseudemoia palfreymani*. Biology of Australian frogs and reptiles (ed. by G. Grigg, R. Shine and H. Hehmann), pp. 477–480. Surrey Beatty & Sons, Sydney.

Rull, V. (2009) Microrefugia. Journal of Biogeography, 36, 481–484.

Sersic, A.N., Cosacov, A., Cocucci, A.A., Johnson, L.A., Pozner, R., Avila, L.J., Sites, J.W. & Morando, M. (2011) Emerging phylogeographical patterns of plants and terrestrial vertebrates from Patagonia. Biological Journal of the Linnean Society, 103, 475–494.

Sérsic, A.N., Cosacov, A., Cocucci, A.A., Johnson, L.A., Pozner, R., Avila, L.J., Sites, J.W. & Morando, M. (2011) Emerging phylogeographical patterns of plants and terrestrial vertebrates from Patagonia. Biological Journal of the Linnean Society, 103, 475–494.

Sheedy, E.M., van de Wouw, A.P., Howlett, B.J. & May, T.W. (2015) Population genetic structure of the ectomycorrhizal fungus *Laccaria* sp A resembles that of its host tree *Nothofagus cunninghamii*. Fungal Ecology, 13, 23–32.

Sigleo, W. & Colhoun, E.A. (1981) A short pollen diagram from Crown Lagoon in the Midlands of Tasmania. Papers and Proceedings of the Royal Society of Tasmania, 115, 181–188.

Sinervo, B., Méndez-de-la-Cruz, F., Miles, D.B., Heulin, B., Bastiaans, E., Villagrán-Santa Cruz, M., Lara-Resendiz, R., Martínez-Méndez, N., Calderón-Espinosa, M.L., Meza-Lázaro, R.N., Gadsden, H., Avila, L.J., Morando, M., De la Riva, I.J., Sepulveda, P.V., Rocha, C.F.D., Ibargüengoytía, N., Puntriano, C.A., Massot, M., Lepetz, V., Oksanen, T.A., Chapple, D.G., Bauer, A.M., Branch, W.R., Clobert, J. & Sites, J.W. (2010) Erosion of Lizard Diversity by Climate Change and Altered Thermal Niches. Science, 328, 894–899.

Soltis, D.E., Morris, A.B., McLachlan, J.S., Manos, P.S. & Soltis, P.S. (2006) Comparative phylogeography of unglaciated eastern North America. Molecular Ecology, 15, 4261–4293.

Stephens, M., Smith, N.J. & Donnelly, P. (2001) A new statistical method for haplotype reconstruction from population data. American Journal of Human Genetics, 68, 978–989.

Stewart, J.R. & Lister, A.M. (2001) Cryptic northern refugia and the origins of the modern biota. Trends in Ecology & Evolution, 16, 608–613.

Stewart, J.R., Lister, A.M., Barnes, I. & Dalen, L. (2010) Refugia revisited: individualistic responses of species in space and time. Proceedings of the Royal Society B-Biological Sciences, 277, 661–671.

Swain, R. & Jones, S.M. (2000a) Maternal effects associated with gestation conditions in a viviparous lizard, *Niveoscincus metallicus*. Herpetological Monographs, 14, 432–440.

Swain, R. & Jones, S.M. (2000b) Facultative placentotrophy: half-way house or strategic solution? Comp Biochem Physiol A Mol Integr Physiol, 127, 441–51.

Symula, R., Keogh, J.S. & Cannatella, D.C. (2008) Ancient phylogeographic divergence in southeastern Australia among populations of the widespread common froglet, *Crinia signifera*. Molecular Phylogenetics and Evolution, 47, 569–580.

Taberlet, P., Fumagalli, L., Wust-Saucy, A.G. & Cosson, J.F. (1998) Comparative phylogeography and postglacial colonization routes in Europe. Molecular Ecology, 7, 453–464.

Tamura, K., Stecher, G., Peterson, D., Filipski, A. & Kumar, S. (2013) MEGA6: Molecular Evolutionary Genetics Analysis Version 6.0. Molecular Biology and Evolution, 30, 2725–2729.

Turchetto-Zolet, A.C., Pinheiro, F., Salgueiro, F. & Palma-Silva, C. (2013) Phylogeographical patterns shed light on evolutionary process in South America. Molecular Ecology, 22, 1193–1213.

Turner, T.F. & Trexler, J.C. (1998) Ecological and historical associations of gene flow in darters (Teleostei: Percidae). Evolution, 52, 1781–1801.

Tzedakis, P.C., Emerson, B.C. & Hewitt, G.M. (2013) Cryptic or mystic? Glacial tree refugia in northern Europe. Trends in Ecology & Evolution, 28, 696–704.

Uller, T., While, G.M., Cadby, C.D., Harts, A., O’Connor, K., Pen, I. & Wapstra, E. (2011) Altitudinal divergence in maternal thermoregulatory behaviour may be driven by differences in selection on offspring survival in a viviparous lizard. Evolution, 65, 2313–2324.

Vera-Escalona, I., D’Elía, G., Gouin, N., Fontanella, F.M., Muñoz-Mendoza, C., Sites, J.W., Jr. & Victoriano, P.F. (2012) Lizards on Ice: Evidence for Multiple Refugia in L*iolaemus pictus* (Liolaemidae) during the Last Glacial Maximum in the Southern Andean Beech Forests. PLoS one, 7, e48358.

Wallis, G.P. & Trewick, S.A. (2009) New Zealand phylogeography: Evolution on a small continent. Molecular Ecology, 18, 3548–3580.

Wallis, G.P., Waters, J.M., Upton, P. & Craw, D. (2016) Transverse Alpine Speciation Driven by Glaciation. Trends in Ecology & Evolution, 31, 916–926.

Wapstra, E., Swain, R., Jones, S.M. & O’Reilly, J. (1999) Geographic and annual variation in reproductive cycles in the Tasmanian spotted snow skink, *Niveoscincus ocellatus* (Squamata: Scincidae). Australian Journal of Zoology, 47, 539–550.

Wapstra, E., Uller, T., Sinn, D.L., Olsson, M., Mazurek, K., Joss, J. & Shine, R. (2009) Climate effects on offspring sex ratio in a viviparous lizard. Journal of Animal Ecology, 78, 84–90.

Waters, J.M. (2011) Competitive exclusion: phylogeography’s ‘elephant in the room’? Molecular Ecology, 20, 4388–4394.

Waters, J.M., Fraser, C.I. & Hewitt, G.M. (2013) Founder takes all: density-dependent processes structure biodiversity. Trends in Ecology & Evolution, 28, 78–85.

Whiteley, A.R., Spruell, P. & Allendorf, F.W. (2004) Ecological and life history characteristics predict population genetic divergence of two salmonids in the same landscape. Molecular Ecology, 13, 3675–3688.

Williams, C.F. (1994) Genetic consequences of seed dispersal in 3 sympatric forest herbs. 2. microspatial genetic structure within populations. Evolution, 48, 1959–1972.

Worth, J.R.P., Jordan, G.J., McKinnon, G.E. & Vaillancourt, R.E. (2009) The major Australian cool temperate rainforest tree *Nothofagus cunninghami*i withstood Pleistocene glacial aridity within multiple regions: evidence from the chloroplast. New Phytologist, 182, 519–532.

Worth, J.R.P., Marthick, J.R., Jordan, G.J. & Vaillancourt, R.E. (2011) Low but structured chloroplast diversity in Atherosperma moschatum (Atherospermataceae) suggests bottlenecks in response to the Pleistocene glacials. Annals of botany, 108, 1247–1256.

Zhang, Z.Y., Cashins, S., Philips, A. & Burridge, C.P. (2014) Significant population genetic structuring but a lack of phylogeographic structuring in the endemic Tasmanian tree frog (Litoria burrowsae). Australian Journal of Zoology, 62, 238–245.

