## Supplementary Tables and Figures for "Phylogeographic parallelism: concordance of patterns in closely related species illuminates underlying mechanisms in the historically glaciated Tasmanian landscape"

**Supplementary Table 1.** *Niveoscincus metallicus* tissue collection locations. Asterisks denote sites with tissue sourced from museums or other institutions. The number of sequences obtained for both ND2 and ND4, for ND4 alone, and for β-globin is listed. Tissue sources: CMC = collected by Colin McCoull; CKJ(ANU) = collected or obtained by Charlotte Jennings at the Australian National University; SAM = South Australia Museum; NMV = National Museum Victoria; ANWC = Australian National Wildlife Collection; MD = Michael Driessen, DPIPWE, Tasmania. Latitudes and longitudes (coordinate system: WGS84) are given in decimal degrees.

| Location | Tissue source | Latitude | Longitude | Tissue samples | ND2+ND4 | ND4 only | β-globin |
| --- | --- | --- | --- | --- | --- | --- | --- |
| Beauty Point | Field | -41.1495 | 146.8180 | 8 | 5 | 5 | 4 |
| Ben Lomond* | Field/SAM | -41.5025 | 147.6135 | 22 | 6 | 15 | - |
| Bicheno | Field | -41.8759 | 148.3108 | 20 | 6 | 10 | 1 |
| Breona* | SAM | -41.7868 | 146.7030 | 3 | 2 | 2 | - |
| Bridport | Field | -41.0025 | 147.3940 | 9 | 8 | 8 | 1 |
| Cataract Gorge | Field | -41.4401 | 147.1266 | 19 | - | 10 | 1 |
| Clarence Lagoon* | CMC | -42.0860 | 146.3215 | 47 | 16 | 16 | 3 |
| Clarke Island | Field | -40.5125 | 148.1263 | 4 | 4 | 4 | - |
| Coles Bay | Field | -42.1223 | 148.3419 | 15 | 7 | 8 | 1 |
| Cradle Mountain* | CKJ(ANU) | -41.6900 | 145.9500 | 3 | 1 | 4 | - |
| Deal Island* | ANWC | -39.4728 | 147.3150 | 4 | 4 | 4 | - |
| Deloraine | Field | -41.5786 | 146.6409 | 11 | 3 | 5 | 1 |
| Devil’s Gullet | Field | -41.6635 | 146.3214 | 11 | 3 | 7 | - |
| Dove Lake* | CKJ(ANU) | -41.6606 | 145.9624 | 10 | 1 | 2 | 1 |
| Falmouth | CMC | -41.503 | 148.2747 | 9 | 7 | 7 | 3 |
| Fortescue Bay | Field | -43.1383 | 147.9567 | 11 | 9 | 9 | - |
| Garden Island | SAM | -43.2600 | 147.1300 | 3 | 3 | 3 | 2 |
| Hartz Peak* | Field/SAM | -43.2213 | 146.7742 | 3 | 3 | 3 | - |
| Hillwood | Field | -41.2186 | 146.9509 | 2 | 2 | 2 | - |
| Hobart | CMC | -42.8944 | 147.2947 | 30 | 4 | 5 | 5 |
| Hogan Island | NMV | -39.2167 | 146.9833 | 5 | - | - | - |
| Killiecrankie | Field | -39.7988 | 147.8605 | 26 | 5 | 5 | 2 |
| King William Creek* | MD | -43.0951 | 146.1555 | 4 | 2 | 2 | - |
| Lagoon of Islands | Field | -42.1129 | 146.9344 | 1 | 1 | 1 | 1 |
| Lake Augusta | Field | -41.8629 | 146.5531 | 18 | 10 | 10 | 2 |
| Lake Chisholm | Field | -41.1348 | 145.0616 | 3 | 3 | 3 | - |
| Lake St Clair | Field | -42.1161 | 146.1786 | 15 | 10 | 13 | 3 |
| Lost Falls | Field | -42.0431 | 147.8915 | 20 | 8 | 9 | 2 |
| McPartlan Pass* | MD | -42.849 | 146.1909 | 4 | 1 | 1 | 1 |
| Miena | Field | -41.9807 | 146.7277 | 10 | 7 | 10 | - |
| Mount Bischoff | CMC | -41.4328 | 145.5236 | 30 | - | 6 | 3 |
| Mt Nelson | Field | -42.9236 | 147.3439 | 20 | 5 | 5 | - |
| Mt Wellington | CMC | -42.885 | 147.2194 | 51 | 5 | 5 | 4 |
| Mt Baw Baw | CMC | -37.8122 | 146.1283 | 13 | 10 | 10 | 5 |
| Mt Chappell Island* | GenBank | -40.2688 | 147.9359 | NA | 1 | 1 | - |
| Mt Field | Field | -42.6815 | 146.7161 | 20 | 6 | 6 | 1 |
| Mt Oakleigh | SAM | -41.8065 | 146.0476 | 3 | 1 | 2 | - |
| Orford | Field/CMC | -42.5563 | 147.8321 | 72 | 5 | 5 | 3 |
| Pine Lake | Field | -41.7414 | 146.7065 | 2 | 2 | 2 | 2 |
| Quamby Bluff | Field | -41.6603 | 146.7226 | 4 | 1 | 3 | - |
| Rodondo Island | DPIPWE | -39.2318 | 146.3846 | 8 | 8 | 8 | 4 |
| Ross | Field | -42.0323 | 147.4909 | 3 | 3 | 3 | - |
| Russell River | Field | -42.9414 | 146.7880 | 20 | 10 | 10 | 3 |
| Scotts Road (Geeveston) | Field | -43.1976 | 146.9570 | 7 | 7 | 7 | 3 |
| Scottsdale | Field | -41.1612 | 147.5071 | 4 | 4 | 4 | 1 |
| Sideling Range | Field | -41.2671 | 147.3967 | 20 | 10 | 10 | - |
| South Cape | CMC | -43.5863 | 146.886 | 25 | 10 | 10 | 1 |
| Southport | Field | -43.4350 | 146.9665 | 14 | 13 | 13 | 7 |
| The Gardens | Field | -41.1808 | 148.2627 | 18 | 7 | 7 | 1 |
| Tooms Lake | Field | -42.2126 | 147.7828 | 22 | 6 | 6 | 5 |
| Tooradin | CMC | -38.2023 | 145.3796 | 10 | 10 | 10 | 1 |
| Trousers Point | Field | -40.2284 | 148.0310 | 5 | 5 | 5 | - |
| Waddamana | Field | -42.1285 | 146.7551 | 22 | 5 | 5 | 1 |
| Whitemark* | Field/AWNC | -40.1023 | 148.0022 | 8 | - | - | - |
| Wilmot | Field | -41.3909 | 146.1168 | 10 | 3 | 9 | 3 |
| Wilsons Promontory* | NMV | -39.0311 | 146.3247 | 6 | 6 | 6 | 2 |
| Wybalenna | Field | -40.0176 | 147.8792 | 12 | - | - | - |
|  |  | **Samples** | | **779** | **284** | **341** | **84** |
|  |  | **Sites** |  | **57** | **52** | **54** | **35** |

**Supplementary Table 2.** *Niveoscincus metallicus* tissue sourced from additional collections. Not all samples are represented in the final dataset, due to amplification or sequencing failure or to species misidentification. Source: CMC = frozen tissue collected by Colin McCoull; CKJ(ANU) = collected by Charlotte Jennings, ANU; SAM = South Australia Museum; NMV = National Museum Victoria; ANWC = Australian National Wildlife Collection; MD = Michael Driessen, DPIPWE, Tasmania.

| Source | Site | Catalogue number | Collection date |
| --- | --- | --- | --- |
| NMV | Hogan Island | D42409 |  |
| NMV | Hogan Island | D42423 |  |
| NMV | Hogan Island | D42440 |  |
| NMV | Hogan Island | D42443 |  |
| NMV | Hogan Island | D42468 |  |
| ANWC | Whitemark | R05132 |  |
| ANWC | Whitemark | R05133 |  |
| ANWC | Deal Island | R05630 |  |
| ANWC | Deal Island | R05631 |  |
| ANWC | Deal Island | R05632 |  |
| ANWC | Deal Island | R05633 |  |
| NMV | Wilsons Promontory | Z21472 |  |
| NMV | Wilsons Promontory | Z21473 |  |
| NMV | Wilsons Promontory | Z21551 |  |
| NMV | Wilsons Promontory | Z21564 |  |
| NMV | Wilsons Promontory | Z21568 |  |
| NMV | Wilsons Promontory | Z21569 |  |
| MD | McPartlan Pass | Pitfall 9D | 20/11/2001 |
| MD | McPartlan Pass | Pitfall 6D | 13/03/2003 |
| MD | McPartlan Pass | Pitfall 7D | 13/03/2003 |
| MD | McPartlan Pass | Pitfall 9A | 13/03/2003 |
| MD | King William Creek | Pitfall 4E | 1/03/2000 |
| MD | King William Creek | Pitfall 3M | 28/03/2003 |
| MD | King William Creek | Pitfall 5F | 23/03/2000 |
| MD | King William Creek | Pitfall 4G | 1/03/2000 |
| SAM | Ben Lomond | ABTC11192 | |
| SAM | Ben Lomond | ABTC11193 | |
| SAM | Breona | ABTC23146 | |
| SAM | Breona | ABTC23147 | |
| SAM | Dove Lake | ABTC23191 | |
| SAM | Dove Lake | ABTC23192 | |
| SAM | Dove Lake | ABTC23193 | |
| SAM | Dove Lake | ABTC23194 | |
| SAM | Dove Lake | ABTC23195 | |
| SAM | Dove Lake | ABTC23196 | |
| SAM | Dove Lake | ABTC23197 | |
| SAM | Dove Lake | ABTC23198 | |
| SAM | Dove Lake | ABTC23199 | |
| SAM | Dove Lake | ABTC23200 | |
| SAM | Hartz Peak | ABTC23139 | |
| SAM | Mt Oakleigh | ABTC23140 | |
| SAM | Mt Oakleigh | ABTC23141 | |
| SAM | Mt Oakleigh | ABTC23132 | |
| SAM | Mt Oakleigh | ABTC23133 | |
| SAM | Mt Oakleigh | ABTC23391 | |
| ANU | Garden Island | ABTC54699 | |
| ANU | Garden Island | ABTC54699 | |
| ANU | Garden Island | ABTC54685 | |
| ANU | Garden Island | ABTC54686 | |
| ANU | Cradle Mtn | CKJ0248 |  |
| ANU | Cradle Mtn | CKJ0251 |  |
| ANU | Cradle Mtn | CKJ0250 |  |

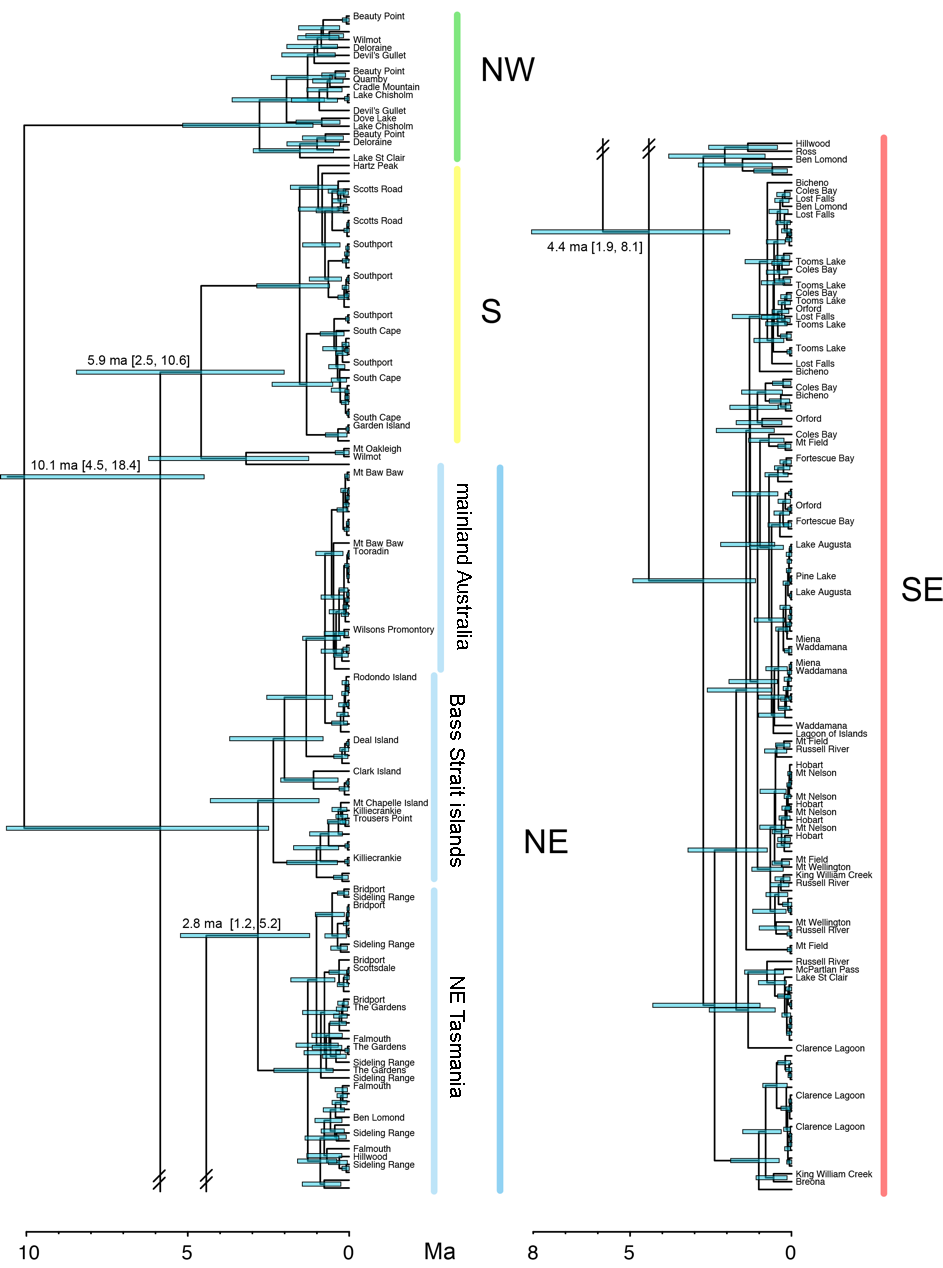

**Supplementary Figure 1.** Maximum clade credibility tree obtained from analysis of 1406 bp of ND2 and ND4 mitochondrial DNA sequences from *Niveoscincus metallicus*. Concatenated mitochondrial sequences were analysed in BEAST2 with a coalescent constant population tree prior and a strict molecular clock with a normally distributed clock rate of 1.52% sequence divergence per million years and a standard deviation of 0.5% divergence. Branch lengths are scaled proportional to time and the scale bar is in millions of years. Node ages are mean branch heights. Horizontal coloured bars on nodes represent 95% High Posterior Density (HPD) of node ages; the node bar for the NW clade has been truncated, and replaced with mean age and upper and lower 95% HPD values. Vertical coloured bars represent major mitochondrial clades and important subdivisions within major clades. The sampling locations of unlabelled tips are the same as the site label immediately above.

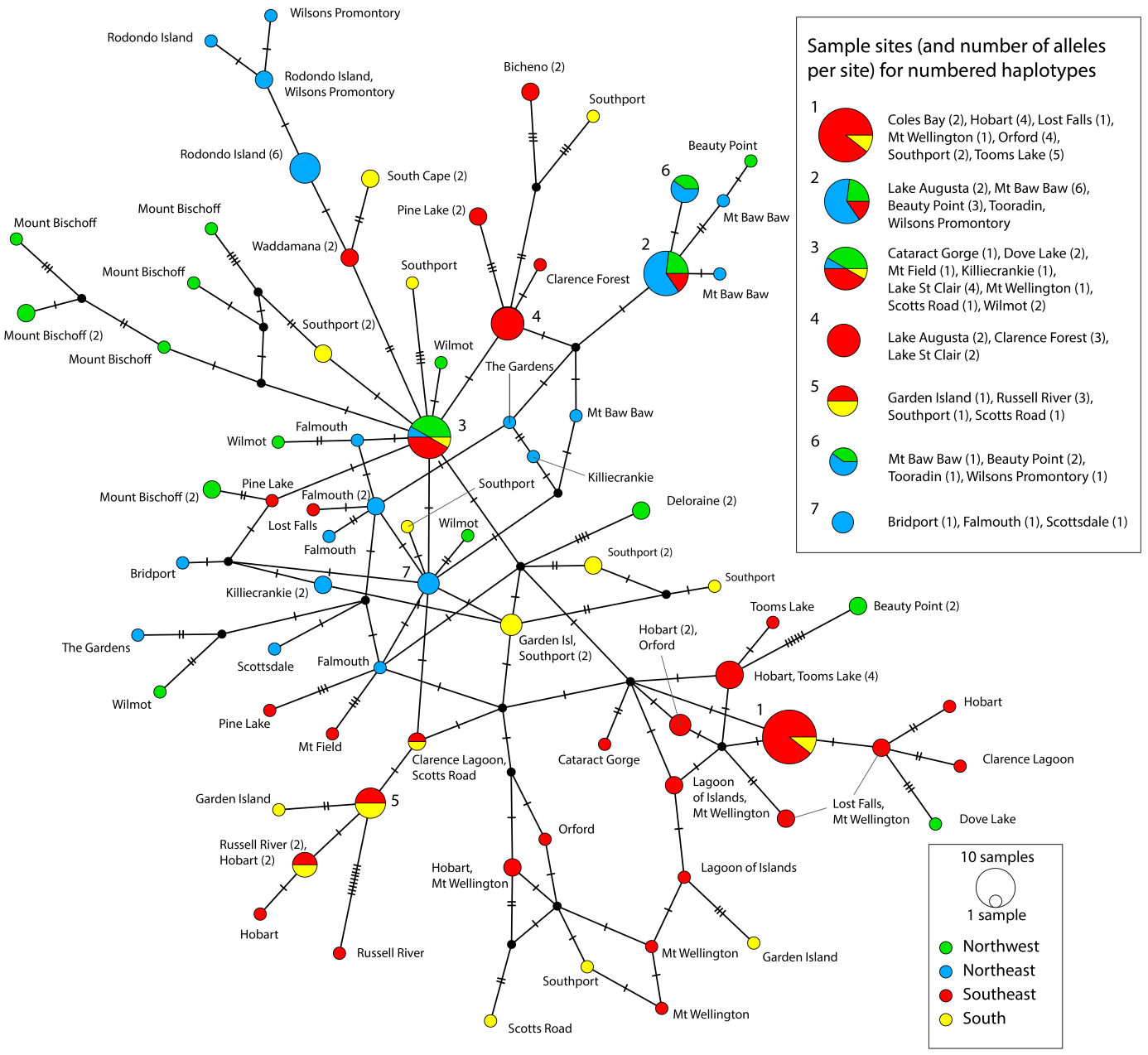

**Supplementary Figure 2.** Network of phased *Niveoscincus metallicus* β-globin alleles constructed using the TCS statistical parsimony algorithm. Coloured circles represent observed haplotypes, and are labelled by sampling site, and colour coded by the mitochondrial clade that each individual was assigned to. The number of alleles per site (if more than one) is indicated in parentheses after the site name. Black circles represent inferred haplotypes. Hatch marks indicate mutations. Circles are scaled proportional to the number of individuals sharing that haplotype. Large circles are numbered and a list of the sites and the number of alleles per site belonging to each numbered haplotype appears at the top right of the diagram.
